# Heteroplasmic Mitochondrial Genetic Variation in the Human Heart

**DOI:** 10.1101/2022.07.13.499890

**Authors:** Liying Xue, Xianbang Sun, Jesse D. Moreira, Yi Li, Karan Kacy Smith, Robert F. Padera, Marc E. Lenburg, Daniel Levy, Emelia J. Benjamin, Deepa M. Gopal, Chunyu Liu, Jessica L. Fetterman

## Abstract

Whether heteroplasmic mitochondrial genetic variants in readily accessible tissues (blood, skeletal muscle) reflect those in the human heart (atrial appendage, left ventricle) is unknown. Using next generation sequencing data from paired tissue samples (n=233) collected postmortem in the Genotype-Tissue Expression project, we identified 558 unique heteroplasmic mitochondrial genetic variants across the four tissues, of which only 13% were shared across all four tissue sites. Between the two cardiac sites, 61% of heteroplasmic mitochondrial genetic variants were unique to one site. A greater proportion of the heteroplasmic variants were non-synonymous or frameshift variants in the muscle sites compared to blood or those variants shared across all four tissues. Compared to blood, the total number of heteroplasmic variants was higher in cardiac tissue, which was associated with advancing age. Our findings suggest that human cardiac tissue has unique heteroplasmic mtDNA variants and may be relevant to aging-related diseases.

## INTRODUCTION

Mitochondrial DNA (mtDNA) is maternally inherited and present in 6-10 copies per mitochondrion (Wallace 2011). Due to the presence of multiple mitochondria per cell, different mtDNAs often co-exist within an individual mitochondrion or population of mitochondria within a cell, a condition termed heteroplasmy. Studies that have evaluated mtDNA variation within populations are restricted to sequencing of the mtDNA in blood samples, which may not reflect mtDNA variation, especially heteroplasmic variation, in other tissues of the same individual (Liu et al. 2021; Laricchia et al. 2022). Pathogenic mtDNA variants undergo purifying selection in hematopoietic cells and hence, may be present in tissues of interest but absent in blood samples from the same individual (Rajasimha et al. 2008). Consistent with this concept, pathogenic mtDNA variants identified in patients with mitochondrial disease are often not detected in blood samples and typically require skeletal muscle biopsy for diagnosis (Rajasimha et al. 2008; Parikh et al. 2015; Stewart and Chinnery 2020).

The accumulation of heteroplasmic variants in mtDNA over time is hypothesized to contribute to an aging-related decline in mitochondrial function and the development of age-related diseases, including cardiovascular disease (Stewart and Chinnery 2015; Wallace 2018). A greater number of heteroplasmic variants in the mtDNA in blood samples is associated with advancing age in the general population and onset and severity of disease in patients with mitochondrial syndromes (Payne et al. 2013; Liu et al. 2018; Wallace 2018). However, whether the alterations in mtDNA identified in blood reflect the mtDNA variants in the human heart or between sites in the heart is not known.

We hypothesized that the presence of heteroplasmic mtDNA variants in different cardiac tissue sites differs from those found in blood samples from the same donor and that advancing age is associated with a greater number of heteroplasmic variants. Using the next generation sequencing available for clinically accessible sites (whole blood, skeletal muscle) and two cardiac sites (atrial appendage, left ventricle), we identified heteroplasmic mtDNA variants that were shared or differed across the tissue sites. We found that heteroplasmic variants differ significantly between the four tissues and even between the cardiac tissues, only 39% of the heteroplasmic variants were shared. Compared to blood, the number of heteroplasmic variants was greater for the cardiac and skeletal muscle tissues, and associated with advancing age.

## RESULTS

We identified mtDNA variants from next generation sequencing (whole-genome sequencing (WGS) and RNA-seq, **Supplemental Fig. S1**) data available through the Genotype-Tissue Expression (GTEx) project from whole blood, atrial appendage, left ventricle, and skeletal muscle collected from the same donors postmortem. Only donors with sequencing data available for all four tissue sites with an average coverage of >400-fold were retained for analysis (**Supplemental Fig. S2**, n=233). Among the donors retained for analysis, 33% were female, 10% were Black, 2% were Asian, and the average age was 54±11 years at the time of death, and comparable to the composition of the entire GTEx sample cohort (**Supplemental Table S1**).

Because RNA-seq was only available for the muscle sites, we compared WGS and RNA-seq in paired blood samples to validate our ability to identify heteroplasmic variants using RNA-seq. We found that the variant allele frequency (VAF) for heteroplasmic variants was highly correlated between WGS and RNA-seq (**Supplemental Fig. S3A**). As the depth of coverage of the mtDNA for blood was greater for WGS compared to RNA-seq (**Supplemental Fig. S3B**), and more similar to the depth of coverage for the muscle sites (**Supplemental Fig. S4**), we used the WGS data for blood for our analyses. The average depth of coverage of the mtDNA from WGS for blood was similar to that of skeletal muscle and slightly lower compared to cardiac tissues (**Supplemental Fig. S4**). The depth for all tissues was sufficient for the detection of low level heteroplasmic variants at VAF >3% (Liu et al. 2021; Wei et al. 2021).

As WGS was performed in blood and RNA-seq with poly-A capture in the muscle sites, the mean depth of coverage was more uniform across the mtDNA for the blood samples (**Supplemental Fig. S5**). We restricted our analyses to loci with an average coverage of 500-fold and we excluded loci in mitochondrial transfer RNAs (tRNAs) due to low coverage. We also excluded low complexity and artifact-prone/known sites (See Methods) (Bar-Yaacov et al. 2013; Wei et al. 2019; Laricchia et al. 2022). The total number of homoplasmic mtDNA variants that were shared across all tissues of an individual donor was similar (**Supplemental Fig. S6**). Only one locus (position 513) had a differing homoplasmic allele across tissues within several donors (n=7) and hence, was deemed artifact and removed from the analyses. We found no correlation of the VAFs of the heteroplasmic variants with postmortem interval (PMI) for any tissue (**Supplemental Fig. S7**).

A total of 558 unique heteroplasmic mtDNA variants were identified with 173 variants unique to blood, 120 unique to heart (atrial appendage, left ventricle), and 65 unique to skeletal muscle (**Fig. 1, Fig. 2A, Supplemental Fig. S8**). Blood, skeletal muscle, and heart only shared 89 heteroplasmic mtDNA variants (16% of all identified heteroplasmic mtDNA variants, **Fig. 2A**), which likely represent maternally inherited variants. The VAFs for the heteroplasmic variants shared across tissues were strongly correlated (**Supplemental Fig. S9**). The overlap in shared heteroplasmic variants between skeletal muscle and heart was similar to the overlap between blood and heart, suggesting that skeletal muscle may not necessarily be a better surrogate tissue than blood for cardiac heteroplasmic variants. Surprisingly, of the 305 variants found in heart, only 120 variants (39%) were shared between the atrial appendage and left ventricle (**Fig. 2B**), indicating that heteroplasmic variants differ across human heart sites.

**Figure 1.**
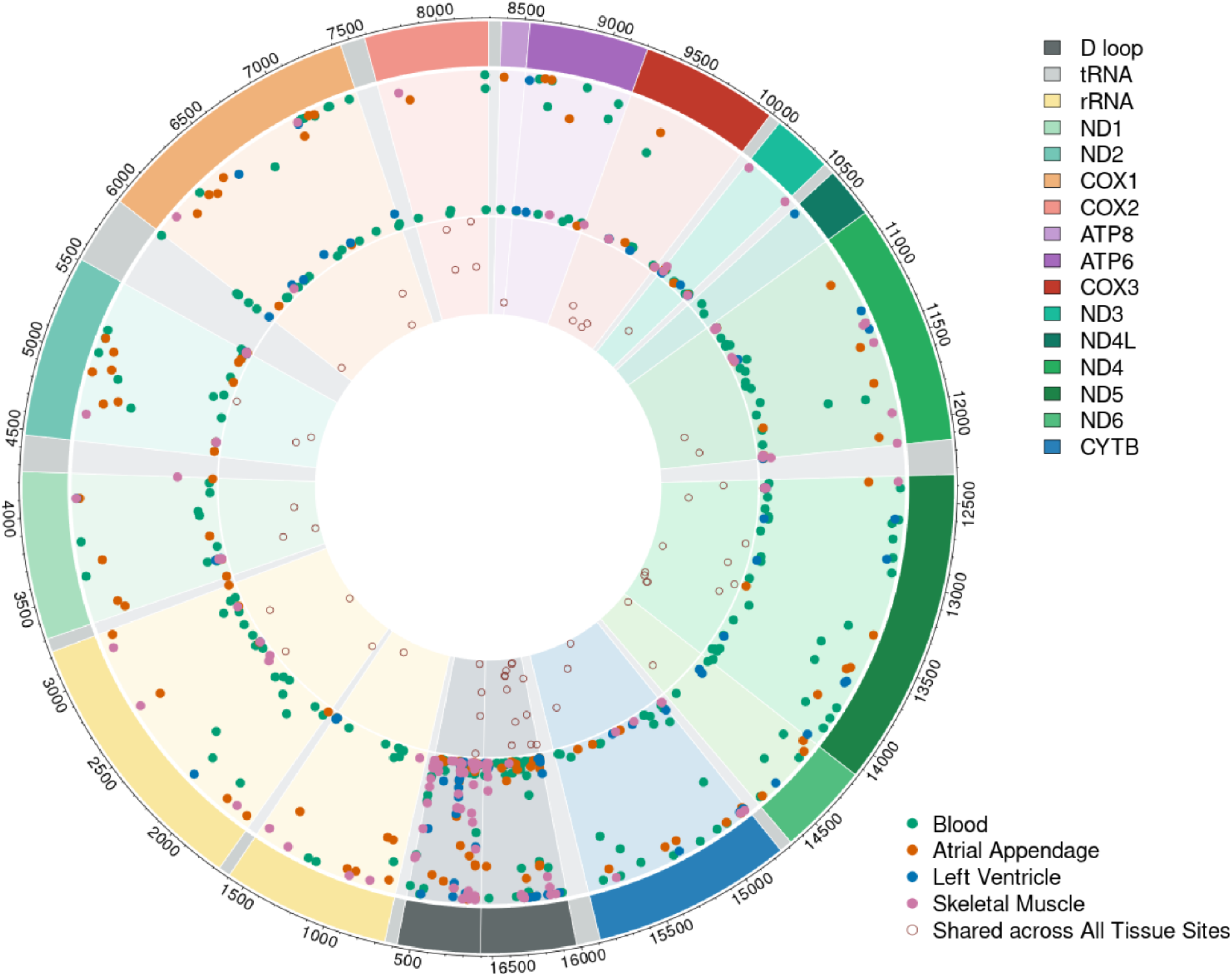
MtDNA Heteroplasmic Variation across Human Tissues. Circos plot of the unique mtDNA heteroplasmic variants identified using WGS and RNA-seq across all 4 tissues with variants found in all tissues indicated in the inner track as open red circles. The middle track indicates the tissue-specific heteroplasmic variants with those in green identified only in blood, orange identified only in atrial appendage, blue found only in the left ventricle, and pink are those identified only in skeletal muscle. Dot height indicates the VAF. The outer track indicates the gene and the position.

**Figure 2.**
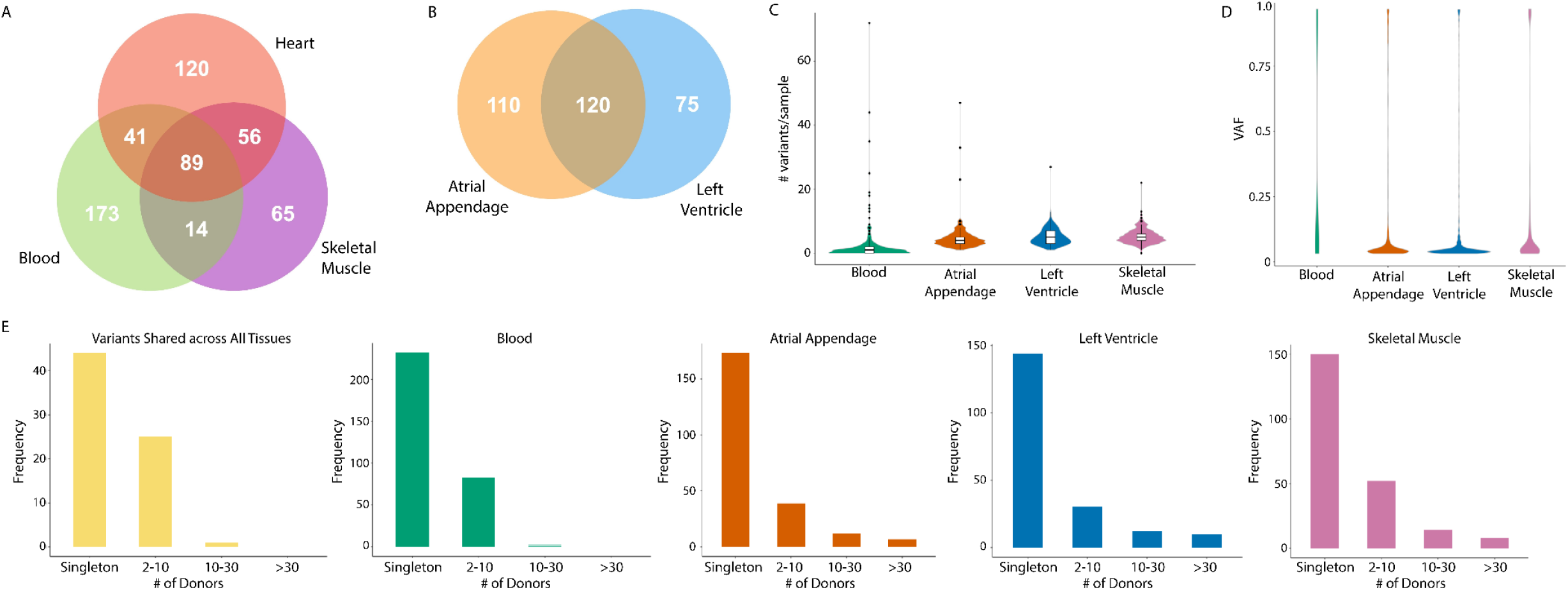
Different Heteroplasmic mtDNA Variants are found in Different Human Tissues. Venn diagram summarizing the number of unique heteroplasmic variants detected in heart, blood, and skeletal muscle (**A**) and between atrial appendage and left ventricle (**B**) across donors. The median number of heteroplasmic variants detected per sample was lower for blood (25-75%, 0-2 variants/donor) compared to the other tissues (blood vs atrial appendage, 25-75%, 3-5 variants/donor, p-value < 2.2 × 10^−16^; blood vs left ventricle, 25-75%, 3-7 variants/donor, p-value < 2.2 × 10^−16^; blood vs skeletal muscle, 25-75%, 4-6 variants/donor, p-value < 2.2 × 10^−16^, (**C**). Most heteroplasmic variants were present at low VAFs (**D**). Most heteroplasmic variants identified were found in a single donor but the muscle sites had a greater proportion of heteroplasmic variants shared among >10 donors (**E**).

All donors had heteroplasmic mtDNA variants in both cardiac sites and 232 of the 233 donors had heteroplasmic variants in skeletal muscle. However, heteroplasmic variants were detected in blood samples from only 62% of donors, which is consistent with prior population studies (Liu et al. 2021; Laricchia et al. 2022). The median number of heteroplasmic variants was lower for blood (median 1 variant/donor) compared to cardiac (atrial appendage, median 4 variants/donor, p < 2.2 × 10^−16^; left ventricle, median 5 variants/donor, p < 2.2 × 10^−16^) and skeletal muscle sites (median 5 variants/donor, p < 2.2 × 10^−16^ **Fig. 2C**). The majority of the heteroplasmic variants were present at VAFs <10% (48-83% of variants, **Fig. 2D**). Most of the heteroplasmic variants identified were found in a single individual (**Fig. 2E**). However, a greater proportion of the heteroplasmic variants identified in the cardiac or skeletal muscle sites were present in >10 donors compared to those found in blood or shared across all tissue sites.

Most heteroplasmic variants identified were transitions (74%, **Fig. 3A**) of a purine to a purine or pyrimidine to a pyrimidine, consistent with prior studies (Greaves et al. 2012; Liu et al. 2018; Laricchia et al. 2022). A greater proportion of the heteroplasmic variants in atrial appendage, left ventricle, and skeletal muscle were frameshift variants compared to blood or those heteroplasmic variants that were shared across all tissues (**Fig. 3B**). In contrast, a greater proportion of the heteroplasmic variants found in blood were synonymous (**Fig. 3B**) and thus, not expected to change the amino acid sequence. Most of the heteroplasmic variants in protein-encoding genes were located in complex I, which is unsurprising considering that 7 of the 13 protein-encoding genes on the mtDNA encode complex I subunits (**Fig. 3C**). The ratio of non-synonymous to synonymous variants was higher for the heteroplasmic variants found only in atrial appendage, left ventricle, and skeletal muscle compared to blood or those variants shared across all tissues (**Fig. 3D**). A greater proportion of the heteroplasmic variants in atrial appendage, left ventricle, and skeletal muscle that were present in >10 donors were frameshift variants (**Fig. 3E**). This suggests that the heteroplasmic variants unique to the cardiac sites or skeletal muscle are more likely to alter the amino acid residue or sequence, which may have implications for the function of the complex.

**Figure 3.**
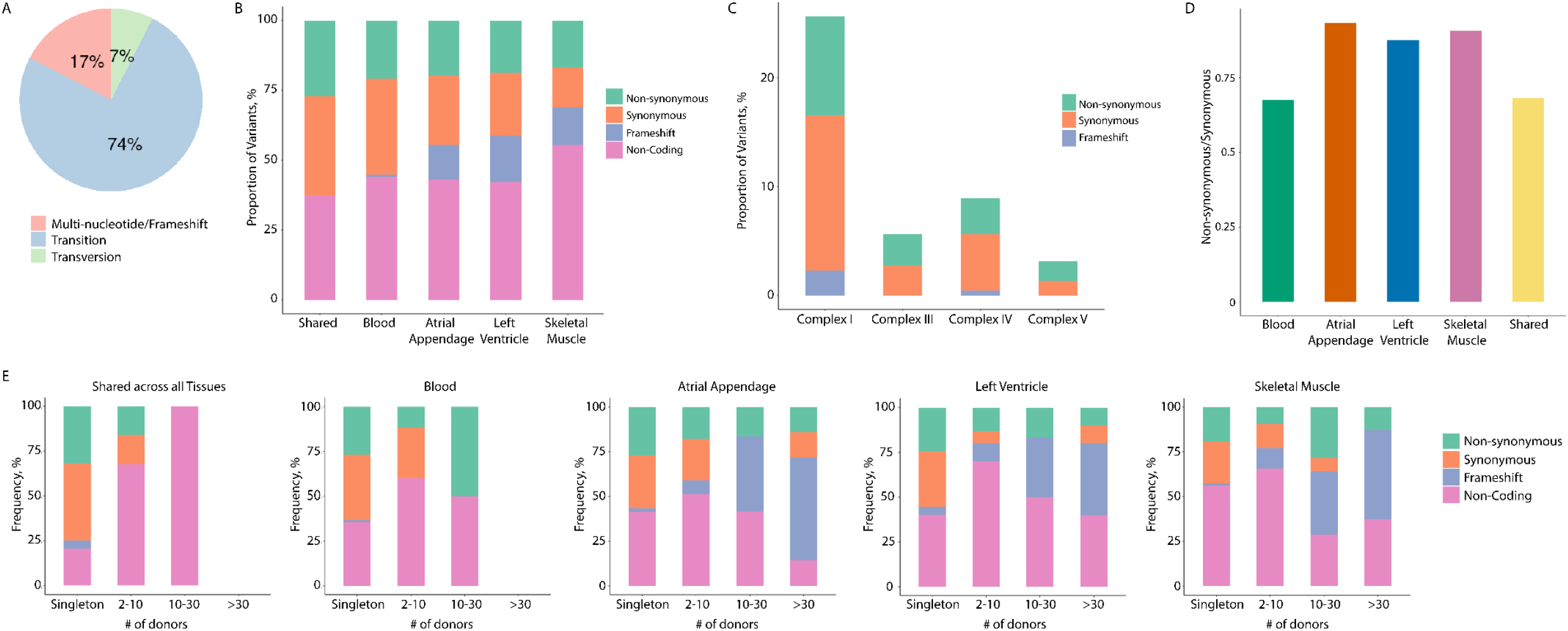
Characterization of the Heteroplasmic mtDNA Variants across the Tissues. Most of the heteroplasmic variants identified were transitions (**A**). Blood had a greater proportion of heteroplasmic variants that were synonymous compared to the muscle sites whereas the muscle sites had a greater proportion of frameshift variants (**B**). Most heteroplasmic variants in the coding region were found in Complex I, which likely reflects the greater number of Complex I subunits encoded by the mtDNA (**C**). The ratio of non-synonymous to synonymous variants was higher for atrial appendage, left ventricle, and skeletal muscle compared to those in blood or shared across all tissues (**D**). A greater proportion of the heteroplasmic variants present in the atrial appendage, left ventricle, or skeletal muscle in >10 donors were frameshift mutations (**E**).

Of the heteroplasmic variants identified, 11 variants were frameshift variants, involving the insertion or deletion of several base pairs so that the reading frame is no longer divisible by three (**Fig. 4**). Most of the frameshift variants were located within complex I encoding genes and were largely restricted to the muscle sites. All frameshift variants had a VAF between 3-10%. With the exception of m.5501A>AG and m.5506TC>T, all other frameshift variants were truncating, in that a stop codon was introduced as a result of the insertion or deletion of nucleotides. Two of the frameshift variants, m.10813C>CA and m.12417C>CA, were recurrent in the atrial appendage, left ventricle, and skeletal muscle of most donors.

**Figure 4.**
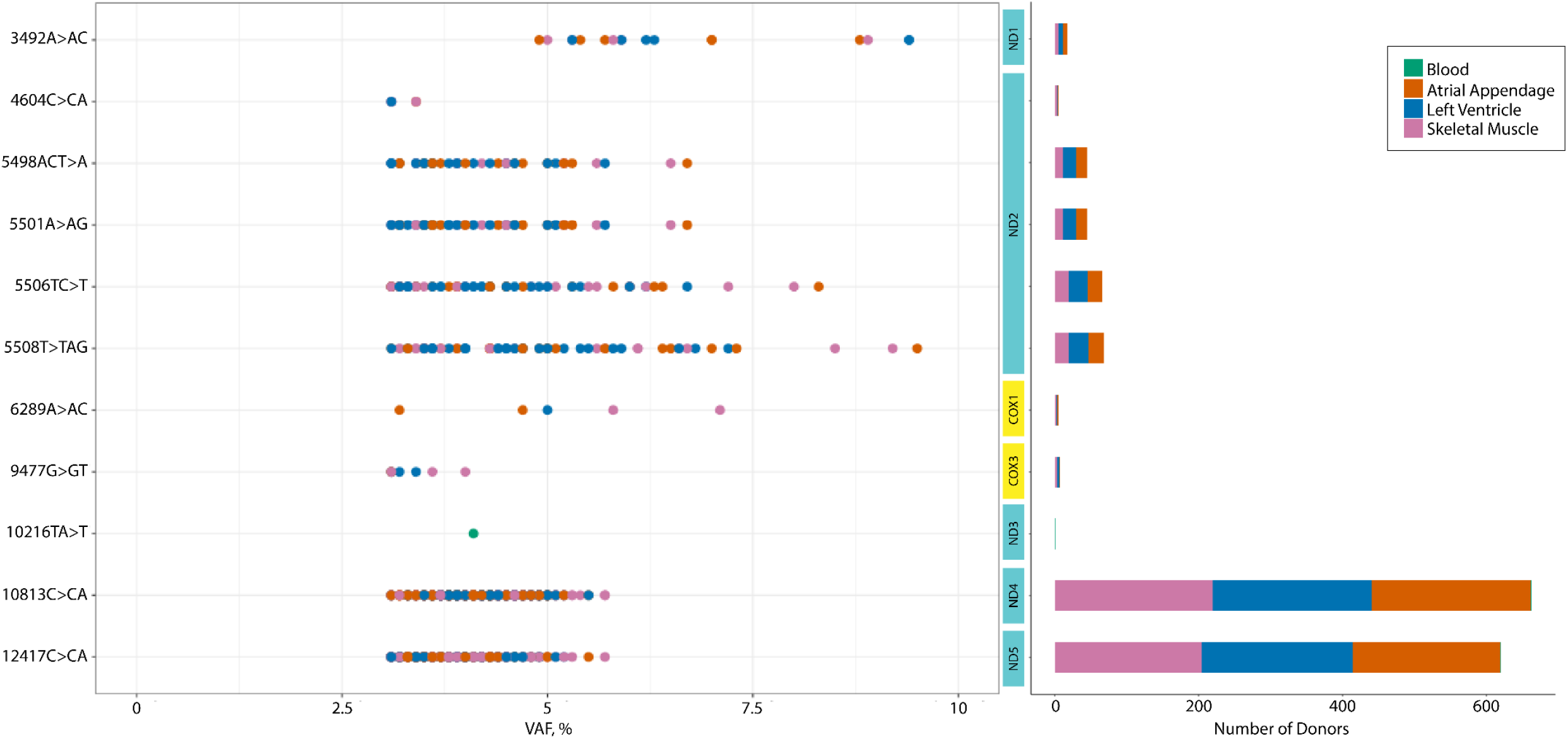
Frequency of Frameshift Variants by Tissue. Several frameshift variants were identified among the heteroplasmic mtDNA variants, primarily in atrial appendage, left ventricle, and skeletal muscle. All frameshift variants identified had VAFs between 3-10% and most were within complex I genes (*ND*). *ND* = NADH dehydrogenase, *COX* = Cytochrome c oxidase

Using Fisher’s exact test, we evaluated whether the tissue specific heteroplasmic variants occurred more or less frequently in the protein-coding genes than would be expected (**Table 1**). Heteroplasmic variants occurred more frequently in *ND2* (OR 1.73, p-value = 0.02) and less frequently than expected in *ND4L* (OR 0.0, p-value = 0.04) in the atrial tissue. No enrichment for heteroplasmic variants in mtDNA genes was observed for the left ventricle or blood, suggesting a random distribution across the protein-encoding genes. For skeletal muscle, heteroplasmic variants occurred less frequently in *COX1* (OR, 0.36, p-value = 0.003) and *ATP6* (OR, 0.24, p-value = 0.03). The differences in frequency of heteroplasmic variants in specific protein-encoding genes across tissues suggests that selective pressures may be operating in a tissue-specific manner.

**Table 1.** Enrichment of Heteroplasmic Variants by Gene.

| **Gene** | **Blood** |  | **Atrial Appendage** |  | **Left Ventricle** |  | **Skeletal Muscle** |  |
| --- | --- | --- | --- | --- | --- | --- | --- | --- |
|  | **OR** | **p-value** | **OR** | **p-value** | **OR** | **p-value** | **OR** | **p-value** |
| *ND1* | 0.67 | 0.21 | 1.15 | 0.55 | 0.69 | 0.41 | 0.69 | 0.44 |
| *ND2* | 0.83 | 0.55 | **1.73** | **0.02** | 1.02 | 0.87 | 1.06 | 0.77 |
| *ND3* | 0.95 | 1.00 | 1.13 | 0.81 | 1.42 | 0.41 | 1.49 | 0.31 |
| *ND4* | 0.74 | 0.24 | 0.60 | 0.10 | 0.54 | 0.09 | 0.91 | 0.90 |
| *ND4L* | 0.36 | 0.18 | **0.00** | **0.04** | 0.32 | 0.38 | 0.00 | 0.05 |
| *ND5* | 1.37 | 0.06 | 0.80 | 0.38 | 1.14 | 0.54 | 0.62 | 0.10 |
| *ND6* | 0.83 | 0.74 | 0.73 | 0.69 | 0.92 | 1.00 | 0.31 | 0.10 |
| *CYB* | 1.23 | 0.30 | 0.88 | 0.78 | 0.75 | 0.54 | 0.65 | 0.25 |
| *COX1* | 0.76 | 0.27 | 0.63 | 0.12 | 0.73 | 0.35 | **0.36** | **0.003** |
| *COX2* | 0.63 | 0.24 | 0.56 | 0.23 | 0.55 | 0.33 | 0.60 | 0.36 |
| *COX3* | 0.47 | 0.05 | 0.78 | 0.62 | 0.86 | 0.86 | 0.74 | 0.61 |
| *ATP6* | 1.05 | 0.77 | 0.79 | 0.73 | 0.70 | 0.56 | **0.24** | **0.03** |
| *ATP8* | 1.06 | 0.79 | 1.13 | 0.75 | 1.42 | 0.47 | 0.40 | 0.52 |

OR = Odds ratio, *ND* = NADH dehydrogenase, *CYB* = Cytochrome b, *COX* = Cytochrome c oxidase, *ATP* = ATP synthase

In MITOMAP, 44 point mutations in protein-encoding genes or non-coding regions are considered confirmed pathogenic mtDNA variants and 538 variants are suspected to be pathogenic (Lott et al. 2013). None of the confirmed pathogenic mtDNA variants were found among the heteroplasmic variants in any of the tissues. However, 43 suspected pathogenic mtDNA variants were identified among the heteroplasmic variants found in blood, atrial appendage, left ventricle, or skeletal muscle at varying VAFs (**Fig. 5**). Most of the suspected pathogenic variants were present in less than 20 donors with the exception of m.185G>A and m.16093T>C in the control region of the mtDNA.

**Figure 5.**
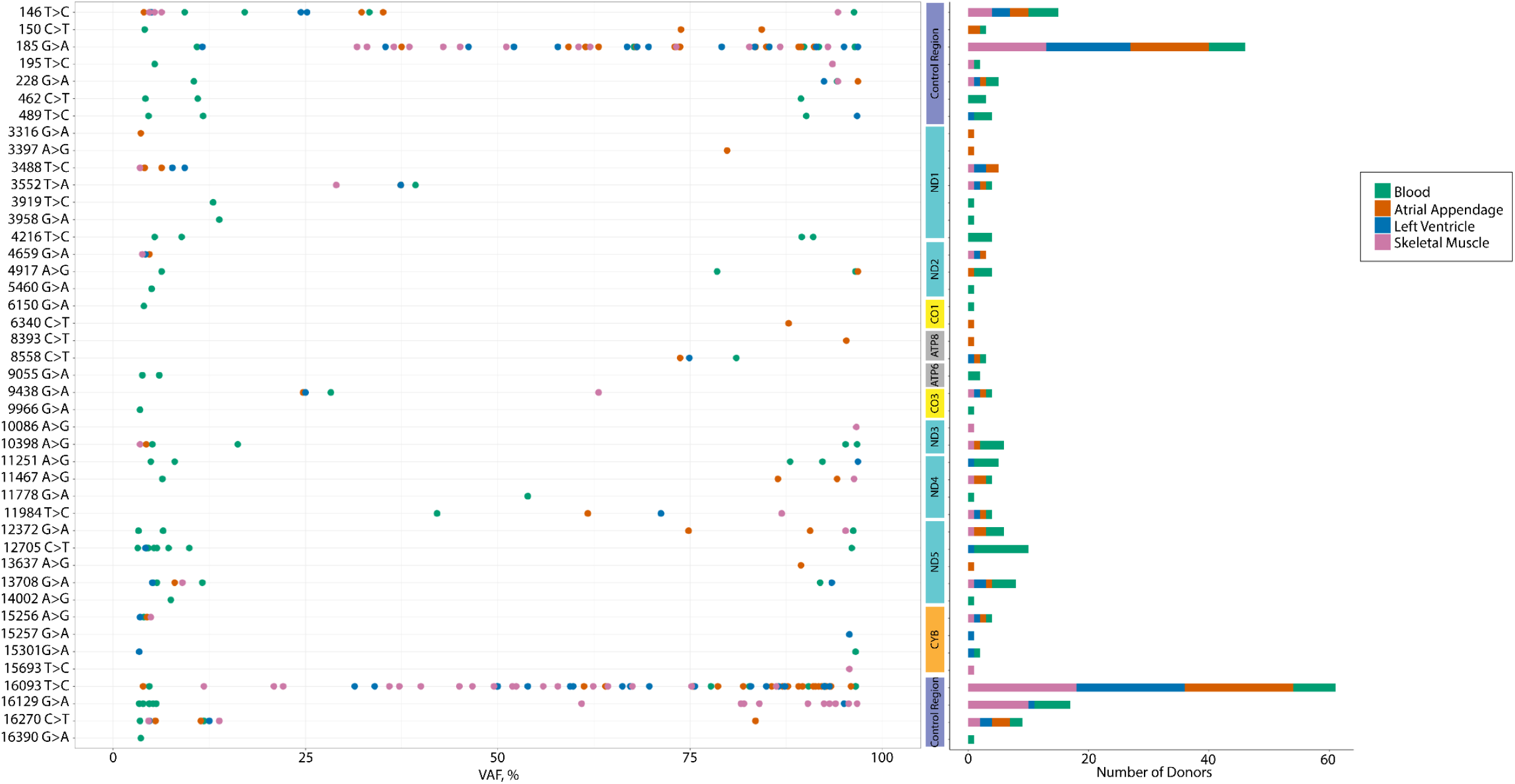
Suspected Pathogenic Mutations Among the Identified Heteroplasmic Variants. Of the 538 suspected pathogenic mtDNA mutations on MITOMAP(Lott et al. 2013), 43 mutations were among the heteroplasmic variants identified across the tissues and at varying VAFs with most variants present in <20 donors. *ND* = NADH dehydrogenase, *CYB* = Cytochrome b, *CO* = Cytochrome c oxidase, *ATP* = ATP synthase

Using linear regression models, we evaluated the association of the number of heteroplasmic variants with donor age in each tissue. We found no correlation of age with the number of heteroplasmic variants in blood (coefficient estimate = −0.71, p-value = 0.4, **Fig. 6**). However, the number of heteroplasmic variants in atrial appendage, left ventricle, and skeletal muscle was positively correlated with age (atrial appendage, coefficient estimate = 2.24, p-value = 0.002; left ventricle, coefficient estimate = 3.19, p-value = 1.2 × 10^−5^; skeletal muscle, coefficient estimate = 2.49, p-value = 0.0007), suggesting that a greater number of heteroplasmic variants in the heart is associated with advancing age.

**Figure 6.**
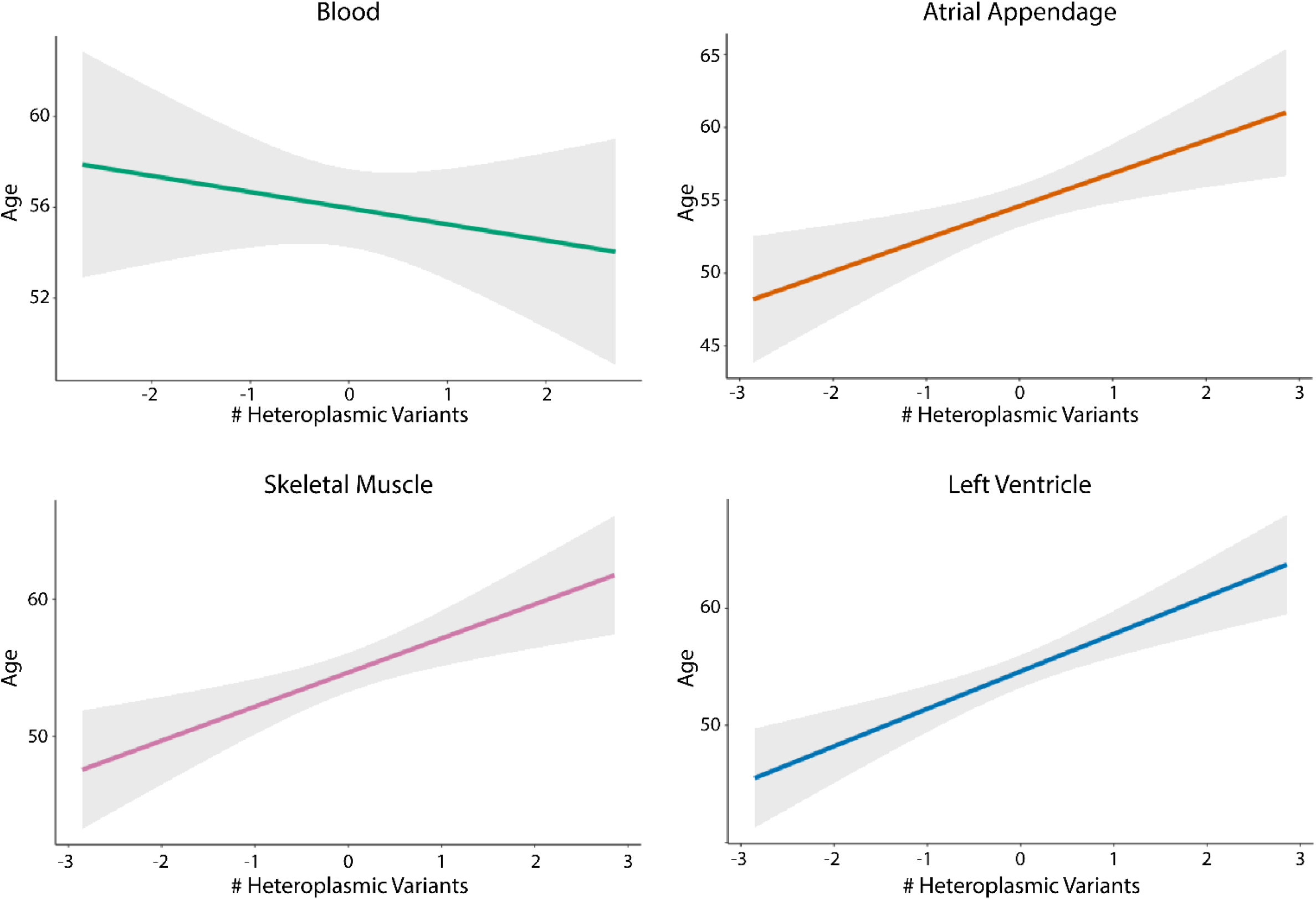
Number of Heteroplasmic Variants in the Cardiac Sites is Associated with Age. Although donor age did not correlate with the number of heteroplasmic variants (inverse normal transformed) in blood (coefficient = −0.71, p-value = 0.4), we found a positive correlation of age with the number of heteroplasmic variants in the atrial appendage (coefficient = 2.24, p-value = 0.002), left ventricle (coefficient = 3.19, p-value =1.2 × 10^−5^), and skeletal muscle (coefficient = 2.49, p-value = 0.0007) in linear regression models. The grey shading indicates the 95% confidence interval.

## DISCUSSION

We evaluated heteroplasmic mitochondrial genetic variation across two clinically accessible sites, blood and skeletal muscle, and two cardiac sites, left ventricle and atrial appendage. We found that very few heteroplasmic mtDNA variants were shared across all four tissues and that heteroplasmic variants differed between the two cardiac sites. The heteroplasmic variants identified in the cardiac tissue and skeletal muscle were more likely to be non-synonymous or frameshift variants and therefore, more likely to alter the function of the protein. The frequency of heteroplasmic variants in specific mitochondrial protein-encoding genes differed across tissues, suggesting that selective pressures may be operating in a tissue-specific manner. The number of heteroplasmic variants was higher in the cardiac and skeletal muscle sites compared to blood, which was associated with advancing age.

The mtDNA has a higher mutation rate than the nuclear genome, which lead to the mitochondrial aging hypothesis (Harman 1972). According to the mitochondrial aging hypothesis, the number and VAF of mutations in the mtDNA accumulate over the lifespan, contributing to the observed age-related decline in mitochondrial function and subsequent disease (Harman 1972). Consistent with this concept, higher mtDNA heteroplasmic variant levels in blood samples are associated with advancing age in the general population (Li et al. 2010; Liu et al. 2018; Liu et al. 2021).

Very few studies have evaluated heteroplasmic mtDNA variation in human tissues other than blood. Large-scale deletions in the mitochondrial genome are found in a number of human tissues, including heart and brain tissue, with a greater fraction of mtDNAs carrying the deletion associated with advancing age (Corral-Debrinski et al. 1992; Cortopassi et al. 1992; Lee et al. 1994). A study that performed next generation sequencing of ten human tissues collected postmortem from two unrelated individuals found that most heteroplasmic variants were shared among the tissues but this study focused solely on the non-coding region (D-Loop) of the mtDNA (Samuels et al. 2013). This study also did not evaluate heteroplasmic variants in cardiac tissue (Samuels et al. 2013).

Another study assessed mtDNA heteroplasmic variation in 12 human tissues collected postmortem from 152 donors; however, the tissue-specific heteroplasmic variants were superficially characterized, only one cardiac site was evaluated, and the focus was on point mutations and the liver (Li et al. 2015). Most of the heteroplasmic variants were found in the control region of the mtDNA whereas in our study, we found a more even distribution of heteroplasmic variants across the mtDNA. The differences in the distribution between the studies could be attributed to differences in the fold coverage of the mtDNA and the result of differences in the VAF used for defining heteroplasmic variants. Similar to our findings that different tissues have differing enrichment for or against heteroplasmic variants in particular mtDNA genes, the liver had a greater proportion of non-synonymous, heteroplasmic variants than would be expected by chance, indicative of positive selection for non-synonymous variants (Li et al. 2015). Overall, prior studies evaluating heteroplasmic mtDNA variants across human tissues have focused on variants in the non-coding region and did not assess heteroplasmic mtDNA variants across regions within a given tissue.

In our study, we found that a greater proportion of the tissue-specific heteroplasmic variants in the cardiac sites were frameshift variants, of which, two of the frameshift variants recurred across most donors. Intriguingly, some tumor types harbor a greater number of frameshift variants, primarily in complex I (Gopal et al. 2018a; Gopal et al. 2018b; Yuan et al. 2020; Gorelick et al. 2021). Colorectal, renal, and thyroid tumors have a greater proportion of mitochondrial heteroplasmic variants that are frameshift variants, including truncating variants, in complex I, which is similar to our findings that most heteroplasmic variants, particularly frameshift variants, in the heart are found in complex I genes (Gopal et al. 2018a; Gopal et al. 2018b; Gorelick et al. 2021). Similar to our findings, a number of the frameshift variants were recurrent in certain tumor types and the total number of heteroplasmic mtDNA variants in tumors correlated with the age of the patient (Gopal et al. 2018a; Yuan et al. 2020; Gorelick et al. 2021). The presence of truncating complex I variants was associated with a greater abundance of genes involved in oxidative phosphorylation and mTOR signaling compared to tumors of the same type lacking truncating variants (Gopal et al. 2018a; Gorelick et al. 2021). Further, a greater VAF for the truncating variants was associated with a greater magnitude of differential expression of oxidative phosphorylation genes (Gorelick et al. 2021).

In renal oncocytomas, frameshift variants in complex I are associated with a higher abundance of transcripts of enzymes involved in glutathione biosynthesis and glutathione metabolites, and lower abundance of enzymes involved in glutathione degradation, suggestive of a protective up-regulation of glutathione against oxidative stress (Gopal et al. 2018a). However, the frameshift variants identified in tumors are present at VAFs much higher than those that we observed in the cardiac or skeletal muscle tissues. Whether the frameshift variants present at the VAFs observed in the cardiac tissue affects mitochondrial and cardiac function will require additional investigation.

Our study has several limitations. A prior study performed ultra-deep sequencing on a non-coding region of the mtDNA and *COX3* and found that most heteroplasmic variants were detected in both whole blood and skeletal muscle from the same individual, but at lower VAFs in blood compared to skeletal muscle (Payne et al. 2013). It is possible that heteroplasmic variants detected in skeletal muscle or heart are present in blood but at VAFs that are below our VAF cutoff of 3% and may require ultra-deep sequencing for detection. Our study is cross-sectional in nature and therefore we are unable to evaluate temporal changes in mtDNA heteroplasmic variation, or establish causality. The GTEx study used poly-A selection for library preparation for RNA-seq; hence, we are unable to evaluate the heteroplasmic variation within the tRNAs and some loci within the D-loop. We are not able to determine whether the mtDNA heteroplasmic variation differs by cell type within a tissue as all sequencing was performed on whole tissue. It is likely that different cell types also have different heteroplasmic variants, which will require single cell sequencing.

Our study lends insight into heteroplasmic mtDNA variation in the human heart. We report that many of the heteroplasmic mtDNA variants found in the heart are not present in blood or skeletal muscle samples from the same individual. Even within the heart, the heteroplasmic mtDNA variants differed between the left ventricle and atrial appendage, suggesting regional differences in mtDNA variation. The differences in frequencies of heteroplasmic mtDNA variants in some of the protein-encoding genes in some tissues but not others suggests that different selective pressures may be operating in a tissue or regional-dependent manner. Although most population studies evaluate the association of heteroplasmic mtDNA variation in blood samples with outcomes of interest, our study suggests that we may be missing potentially important mtDNA variants by not looking in the heart tissue itself.

## METHODS

### Dataset and Data Access

WGS data for whole blood samples and RNA-seq data for whole blood, skeletal muscle, left ventricle, and atrial appendage samples from the GTEx project v8 (dbGaP accession phs000424.v8.p2) were downloaded through the AnVIL portal (https://gen3.theanvil.io/). Only RNA-seq data is currently available for skeletal muscle, left ventricle, and atrial appendage sites. Of the 826 donors in the GTEx dataset, 298 donors had sequencing data available for all four tissue sites. Donors that did not have WGS for whole blood samples or did not have RNA-seq for one or more muscle sites were excluded. Following quality control procedures, 235 individuals satisfied the inclusion criteria and were used in the analysis (**Supplemental Fig. S2**).

### Extracting Mitochondrial Sequences, and Identification and Quality Control Procedures for mtDNA Variants

SAMtools was used to extract reads that aligned to the mtDNA (chrM) from the sequencing files (WGS and RNA-seq for blood, RNA-seq for muscle sites, **Supplemental Fig. S1**). Extracted BAM files were converted to FastQ files using SAMtools bam2fq paired-end mode in preparation for sequence alignment. The sequencing reads were aligned to the revised Cambridge Reference Sequence (rCRS) (Andrews et al. 1999) using BWA-MEM for WGS and STAR aligner version 2.7.9a for RNA-seq (Dobin et al. 2013). The aligned BAM files were fixed by GATK for missing read groups and then sorted and indexed by SAMtools. MtDNA variants were identified from the sorted and indexed files using GATK Mutect2 mitochondrial mode (Laricchia et al. 2022). Our software pipeline is available on GitHub-GitHub-ariel208x/mtDNA_heart_variants.

We excluded 2 donors that had an average coverage <400-fold and restricted our analyses to loci with an average depth >500-fold. We excluded the loci within tRNAs due to low coverage, variants in artifact-prone and low-complexity sites for WGS (66-71, 301, 302, 310, 316, 3107, 12418-12425, 16182-16194), and known artifacts specific to RNA-seq (295 C>T, 13710 A>G, and 2617 A>G) (Bar-Yaacov et al. 2013; Wei et al. 2019; Laricchia et al. 2022). Further, we identified an additional six “variants” present in RNA-seq but not WGS of blood samples from the same donors, which we excluded from our analyses as these loci likely represent artifacts (2487, 2617, 6419, 10277, 10306, 13710).

Variants with a VAF of 3-97% were defined as heteroplasmic variants while variants with a VAF > 97% were defined as homoplasmic variants. We identified 1056 unique homoplasmic variants. Homoplasmic variants were compared across tissue sites within each donor to check for discordance. Due to discordance in homoplasmic variants between tissues for multiple samples (7 donors) at position 513, we excluded this site from our analyses. Other than position 513, the homoplasmic variants were concordant across all tissues of the same donor, regardless of the sequencing technology used.

### Statistical Analyses

Tissue specific variants were identified as unique variants present in only one of the four tissues within each donor and shared variants were heteroplasmic variants found in all four tissues of the donor. We used a Wilcoxon rank sum test to perform pairwise comparisons between tissues within a donor. Fisher’s exact test was used to evaluate whether heteroplasmic variants were under- or over-represented within the protein-encoding genes more than would be expected by chance.

We performed a linear regression to evaluate the association of age with the number of heteroplasmic variants in a given tissue. In order to transform the number of heteroplasmic variants (count) to meet the normality assumption, we applied inverse normal transformation (INT) to log(count+1) and used it as the predictive variable to test the correlation between the number of heteroplasmic variants with age (Model 1).

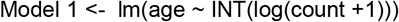

Similar linear regression models were also performed to test for a correlation between the heteroplasmic VAF for the variants in each tissue with PMI and to compare the heteroplasmic VAF for the variants shared across all tissue sites. In Model 2, log transformed PMI was used as the predictor and each heteroplasmic VAF was the outcome.

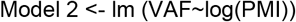

All statistical analyses were performed using R version 4.0.5.

## Acknowledgments

The Genotype-Tissue Expression (GTEx) Project was supported by the Common Fund of the Office of the Director of the National Institutes of Health (commonfund.nih.gov/GTEx). Additional funds were provided by the NCI, NHGRI, NHLBI, NIDA, NIMH, and NINDS. Donors were enrolled at Biospecimen Source Sites funded by NCI\Leidos Biomedical Research, Inc. subcontracts to the National Disease Research Interchange (10XS170), Roswell Park Cancer Institute (10XS171), and Science Care, Inc. (X10S172). The Laboratory, Data Analysis, and Coordinating Center (LDACC) was funded through a contract (HHSN268201000029C) to The Broad Institute, Inc. Biorepository operations were funded through a Leidos Biomedical Research, Inc. subcontract to Van Andel Research Institute (10ST1035). Additional data repository and project management were provided by Leidos Biomedical Research, Inc.(HHSN261200800001E). The Brain Bank was supported supplements to University of Miami grant DA006227. Statistical Methods development grants were made to the University of Geneva (MH090941 & MH101814), the University of Chicago (MH090951,MH090937, MH101825, & MH101820), the University of North Carolina - Chapel Hill (MH090936), North Carolina State University (MH101819),Harvard University (MH090948), Stanford University (MH101782), Washington University (MH101810), and to the University of Pennsylvania (MH101822). The datasets used for the analyses described in this manuscript were obtained from dbGaP at http://www.ncbi.nlm.nih.gov/gap through dbGaP accession number phs000424.v8.p2.

## Sources of Funding

National Heart, Lung, and Blood Institute R21HL144877 (XS, CL)

National Institute on Aging R01AG059727 (XS, CL)

National Heart, Lung, and Blood Institute T32 HL007224-45 (JDM)

National Center for Advancing Translational Sciences, UL1 TR001430 (MEL)

National Heart, Lung, and Blood Institute R01HL092577; American Heart Association AF AHA_18SFRN34110082 (EJB)

American Heart Association grant #17FTF33670369 (DMG)

National Heart, Lung, and Blood Institute K01 HL143142 (JLF)

## Author Contributions

Conceptualization: JLF

Methodology: LX, YL, XS, MEL, DL, EJB, CL, JLF

Investigation: LX, JLF

Visualization: LX, KKS, JDM, MEL, DL, EJB, DMG, CL, JLF

Funding acquisition: JLF

Project administration: JLF

Supervision: CL, DMG, JLF

Writing – original draft: LX, JLF

Writing – review & editing: LX, YL, XS, JDM, KKS, RFP, MEL, DL, EJB, DMG, CL, JLF

